# Olfactory receptors are required for social behavior and neural plasticity in ants, as evidenced by CRISPR-mediated gene knockout

**DOI:** 10.1101/142232

**Authors:** Hua Yan, Comzit Opachaloemphan, Giacomo Mancini, Huan Yang, Matthew Gallitto, Jakub Mlejnek, Kevin Haight, Majid Ghaninia, Lucy Huo, Alexandra Leibholz, Jesse Slone, Xiaofan Zhou, Maria Traficante, Clint A. Penick, Kelly Dolezal, Kaustubh Gokhale, Kelsey Stevens, Ingrid Fetter-Pruneda, Roberto Bonasio, Laurence J. Zwiebel, Shelley Berger, Jüergen Liebig, Danny Reinberg, Claude Desplan

## Abstract

The chemosensory system is key to establishing and maintaining social structure in eusocial insects. Ants exhibit cooperative colonial behaviors reflective of an advanced form of sociality with an extensive dependency on communication. Cuticular hydrocarbons (CHCs) serve as pheromones and cues that regulate multiple aspects of social interactions and behaviors in ants. The perception of CHCs entails odorant receptor neurons (ORNs) that express specific odorant receptors (ORs) encoded by a dramatically expanded *Or* gene family in ants. Until recently, studies of the biological functions of ORs in eusocial insects were stymied by the lack of genetic tools. In most eusocial insect species, only one or a few queens in a colony can transmit the genetic information to their progeny. In contrast, any worker in the ant *Harpegnathos saltator* can be converted into a gamergate (pseudo-queen), and used as a foundress to engender an entire new colony and be crossed for genetic experiments. This feature facilitated CRISPR-Cas9 gene targeting to generate a germline mutation in the *orco* gene that encodes the obligate co-receptor whose mutation should significantly impact ant olfaction. Our results show that Orco exhibits a conserved role in the perception of general odorants but also a role in reproductive physiology and social behavior plasticity in ants. Surprisingly, and in contrast to other insect systems, the loss of OR functionality also dramatically reduces the development of the ant antennal lobe where ORNs project. Taken together, these findings open the possibility of studying the genetics of eusociality and provide inroads towards understanding the function of the expanded ORs family in eusocial insects in regulating caste determination, social communication and neuronal plasticity.

## Introduction

Amongst the most fascinating and enigmatic phenomena in biology is the process by which environmental stimuli drive animal behavior through sensory neuron perception. Eusocial animals, including ants, live in highly sophisticated societies and display a rich repertoire of social behaviors due to the striking division of labor among morphologically and physiologically distinct castes. Most ant castes are established during development in diploid females, while normally, the only role for haploid males is reproduction (Hölldobler and Wilson, 1990). However, the ponerine ant species *Harpegnathos saltator* displays strong adult phenotypic plasticity. When a queen dies or is removed from a colony, workers compete via antennal dueling, and the most frequent duelers become reproductive gamergates (pseudoqueens) (Bonasio, 2012; Hölldobler and Wilson, 2008; Peeters et al., 2000; Sasaki et al., 2016). While these gamergates are not morphologically different from workers, they exhibit queen-like physiology and behavior (e.*g. egg*-laying). This plasticity is likely induced by the deprivation of queen pheromones that are normally communicated via olfactory processes to repress the gamergate transition (Fig. 1A). Indeed, olfaction regulates multiple behaviors in eusocial insects (Ozaki et al., 2005; Sharma et al., 2015), with communication between ant individuals being mediated in part by Cuticular HydroCarbons (CHCs) and other pheromones such as sex, alarm, or trail pheromones (Blomquist et al., 2010; Van Oystaeyen et al., 2014). In *Harpegnathos*, CHC profiles display a clear shift to longer chain hydrocarbons when workers become reproductive gamergates (Liebig et al., 2000). Such a defined variation suggests that queen and gamergate CHCs may serve as pheromones to suppress worker ovary growth and regulate worker-specific behaviors (Sharma et al., 2015; Van Oystaeyen et al., 2014). In *Camponotus* ants, the different profiles of CHCs from nestmates *vs*. non-nestmates are detected by odorant receptor neurons (ORNs), which induce aggression against non-nestmates (Ozaki et al., 2005; Sharma et al., 2015). Although important, these observations remain correlative and genetic tests for the role of olfaction in eusociality are still lacking.

**Figure 1.**
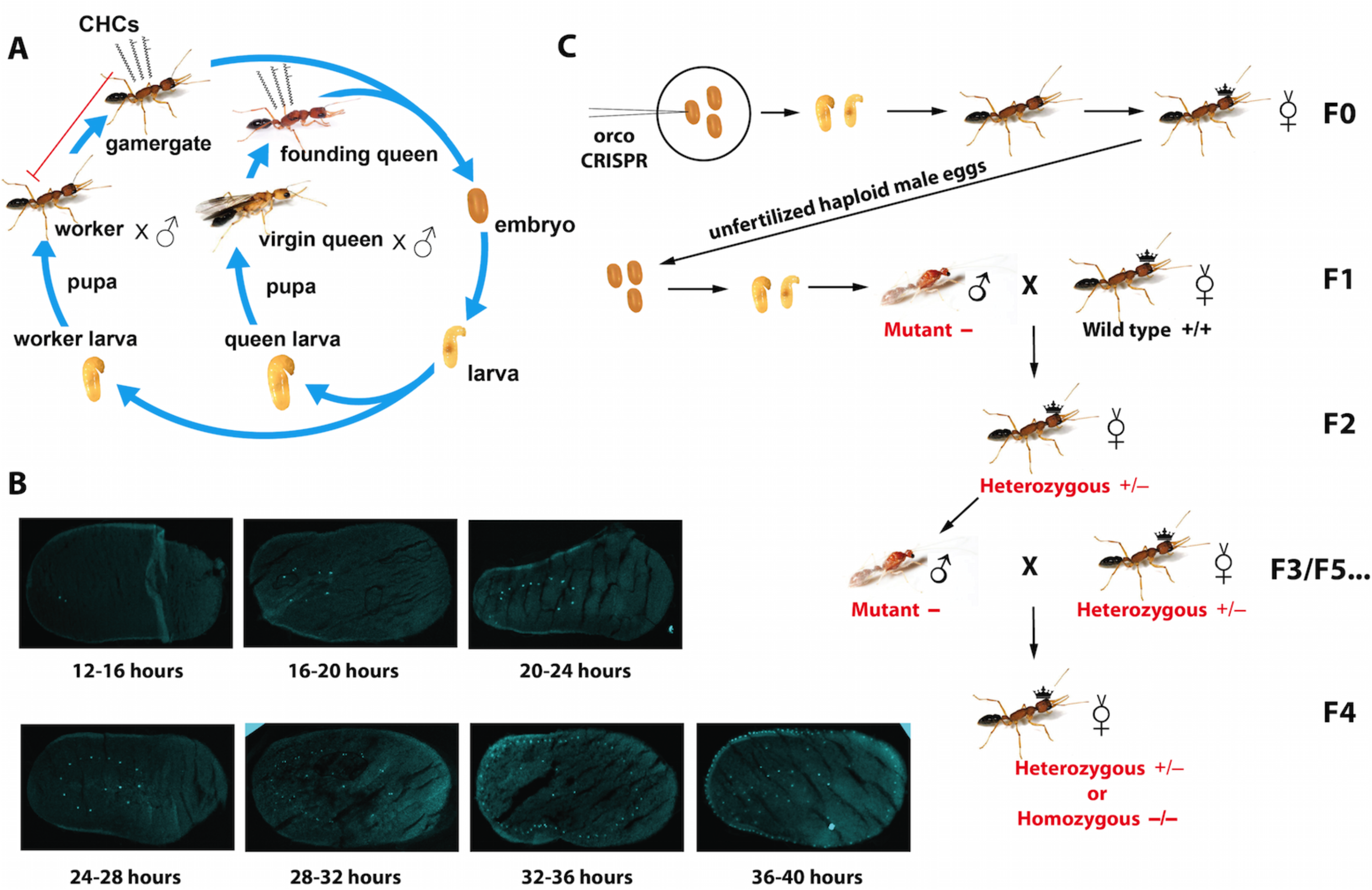
Harpegnathos life cycle, early embryo development, and propagation of the mutant allele. (A) The female embryos develop into larvae. At the 4^th^ instar (Penick et al., 2012), the larvae differentiate into queen vs. worker larvae. The former develop into the callow virgin queens with wings that can found a new colony, while the latter become workers in the nest and, when isolated in group or individually, can become gamergates and lay eggs. Certain cuticular hydrocarbons (CHCs) of queen or gamergate act as a queen pheromone that inhibits worker reproduction. B) Cryosections and DAPI staining of young embryos harvested at different time points after egg deposition (AED, hours below each image) (C) The steps to propagate mutant ants to develop hemizygous males (F3/F5) and homozygous females (F4) for phenotypic analyses. Of note, the F3 mutant males mated with F2 heterozygotes to generate F4 ants, which were half homozygotes and half heterozygotes. The F4 heterozygotes, like F2s, also produced male mutants (F5).

The olfactory sensory system in insects comprises sensory neurons that are located in the sensilla on the antennae, maxillary palps and other sensory appendages (Laissue and Vosshall, 2008). The ORNs express odorant receptors (tuning ORs) that confer specificity to odorant ligands. The repertoire of tuning ORs is encoded by 60 genes in the *Drosophila melanogaster* genome (47 of which are expressed in adults) (Laissue and Vosshall, 2008). However, the family of *Or* genes is dramatically amplified in ants with over 300 *Or* genes having been identified in several ant genomes. The most dramatically expanded *Or* family is the female-specific 9-exon subfamily that has been associated with the evolution of eusocial interactions (Zhou et al., 2015; Zhou et al., 2012). Other insect chemosensory receptor gene families encoding gustatory receptors (GRs) and ionotropic glutamate receptors (IRs) have not undergone a similar expansion and remain in small numbers (Zhou et al., 2015; Zhou et al., 2012). ORN axons project to the antennal lobe (AL) that consists of numerous globule-shaped neuropils known as glomeruli, where the initial synaptic integration occurs before olfactory information is sent via projection neurons to the mushroom body and the lateral horn in the central brain (Wicher, 2015). The *Drosophila* AL contains only 42 glomeruli, while over 400 glomeruli were identified in *Camponotus floridanus* (Zube and Rössler, 2008) and in *Ooceraea biroi* (McKenzie et al., 2016), with more in females than in males (Hoyer et al., 2005; McKenzie et al., 2016). Indeed, in *Ooceraea*, the female-specific 9-exon ORNs project to a large cluster of glomeruli (T6), which are over-represented in female ALs compared to males in both *Ooceraea* and *Camponotus* (McKenzie et al., 2016; Zube and Rössler, 2008).

Orco is the highly conserved pan-olfactory co-receptor in insects that forms an obligate heterodimer with all tuning ORs (Nakagawa et al., 2012; Vosshall and Hansson, 2011; Zhou et al., 2012). The OR-Orco dimers function as ligand-gated ion channels that activate the ORNs upon odorant binding (Benton et al., 2006). Therefore, loss of function in the *orco* gene leads to the loss of all OR-mediated chemosensation. In *Drosophila* and other insects (Larsson et al., 2004)((DeGennaro et al., 2013; Koutroumpa et al., 2016; Li et al., 2016; Yang et al., 2016), *orco* is a non-essential gene whose disruption leads to dramatic reductions in olfactory sensitivity. Mutations in *orco* abolish the behavioral and electrophysiological responses to a number of general odorants in *Drosophila* (Larsson et al., 2004). Loss of *orco* disrupts the female mosquito response to human scent in the absence, but not in the presence of CO_2_, suggesting that other pathways (*e.g*. those mediated by ionotropic olfactory receptors, IRs) are also able to detect odorants (DeGennaro et al., 2013).

In mammals, a very large family of *Or* genes (>1,000) encode G-protein coupled receptors (GPCRs) with no evolutionary relationship to insect ORs. However, as in insects, ORNs only express one OR and all ORNs expressing a given OR project to the same glomeruli in the olfactory bulb. Given the very large number of ORs in mammals, their activity is believed to be required for the ORNs to find their correct glomeruli in the olfactory bulb (Mombaerts, 2006; Yu et al., 2004). In *Drosophila*, with only 42 glomeruli, the presence of an OR is not required for ORNs to project to their correct glomerulus. Indeed, *Drosophila orco* mutants that lack ORN activity do not exhibit altered glomerulus number or morphology (Chiang et al., 2009; Larsson et al., 2004). This suggests that, unlike in mammals, the *Drosophila* brain is hardwired by genetic programs irrespective of environmental stimuli. It is not known whether this hardwiring feature is specific for *Drosophila* or can be generalized to other insect species, such as ants.

Functional studies in ants rest on the ability to genetically manipulate pathways by mutagenesis and transgenesis. Unlike most eusocial insects wherein only the queen in the colony can transmit the genetic information to the next generation, any worker in *Harpegnathos* can be converted to a reproductive gamergate and start a whole new colony (Liebig et al., 1998). This unique feature of this eusocial insect, along with its high-quality genome sequence (Bonasio et al., 2010), provided a strong platform for attempting to generate, propagate and analyze mutants. As ants live in a highly organized social environment, and pheromone and chemical cue detection is thought to be required for proper social communication and colony stability, we targeted the *Harpegnathos orco* gene for mutagenesis. We used the clustered regulatory interspaced short palindromic repeat (CRISPR)/Cas9 system (Cong et al., 2013; Gratz et al., 2013; Hsu et al., 2014), and successfully generated the first germline mutant in ants, which carries a disruption of the *orco* gene. We then generated homozygous females and hemizygous male mutants. Mutation in *orco* not only impaired the ability of the ant to respond to general odorants, as in other insects, but also led to pronounced physiological and behavioral phenotypes. The mutant workers, unlike their age-matched wild type (WT) counterparts, wandered outside of the nests and did not forage to bring cricket preys back to the colony. When isolated, the mutants displayed abnormal behavior reminiscent of dueling during gamergate transition. Yet, they exhibit reduced fertility, laying few eggs and not caring for them. When in a queen-less colony, *orco* mutants did not duel and do not become gamergates, demonstrating that caste determination and reproduction is strongly disrupted by loss of olfaction through ORs. Moreover, *orco* mutant ants also manifested dramatic morphological changes in the AL, suggesting that, in contrast to *Drosophila*, OR activity is essential in establishing proper neural anatomy in ants.

## Results

### Generation of orco KO ants using CRISPR

To maximize the chances for successful genome editing with CRISPR, we sought the best stage (developmental time) at which to inject *Harpegnathos* embryos so that small guide RNAs (sgRNAs) and Cas9 protein would have access to as many nuclei as possible in future germline cells. The development of eusocial insects is relatively longer compared to solitary insects such as *Drosophila* (Yan et al., 2014). In *Harpegnathos saltator*, the time between oviposition to imago eclosion takes 75 days at 25°C, with embryonic development taking 29 days (Fig. 1A). The ideal time for genome editing by microinjection is the syncytial stage, when nuclei divide without cytokinesis (Henderson, 2004), but the timing of this phase in *Harpegnathos* was unknown. We visualized embryos during early embryonic development and found that they reached gastrulation ~48 hours after egg deposition (AED) (Fig. S1A). Nuclear staining of early embryo sections revealed that the syncytial phase lasted from 12 to 36 hours AED (Fig.1B). Based on these observations, we chose this time range to deliver sgRNAs and Cas9 by lateral microinjections (Watanabe et al., 2014). After injection, the embryos were maintained on agar plates until the first hatching. Freshly hatched larvae were transferred to a small WT colony to be reared into pupae and adults by nursing workers (Fig. S1B,C).

Orco is a 7-transmembrane domain protein, with its N-terminus located at the intracellular space (Fig. 2C). We used Cas9 protein (PNA Bio Inc.) and single or multiple *in vitro* synthesized sgRNAs to target the first exon of the *Harpegnathos orco* gene, 50 nucleotides downstream of the translational initiation codon (ATG; Fig. 2A). The target was validated after extracting genomic DNA from injected embryos followed by PCR and pGEM cloning, which identified multiple short deletions from a single sgRNA (Fig. 2A). Longer deletions between the two target sites were found when two sgRNAs were coinjected (Fig. S2A). The efficiency of somatic mutagenesis reached approximately 40%, as evidenced by MiSeq deep sequencing (Kistler et al., 2015) of genomic DNA derived from whole embryos sacrificed 3 days after injections (Fig. S2B).

**Figure 2.**
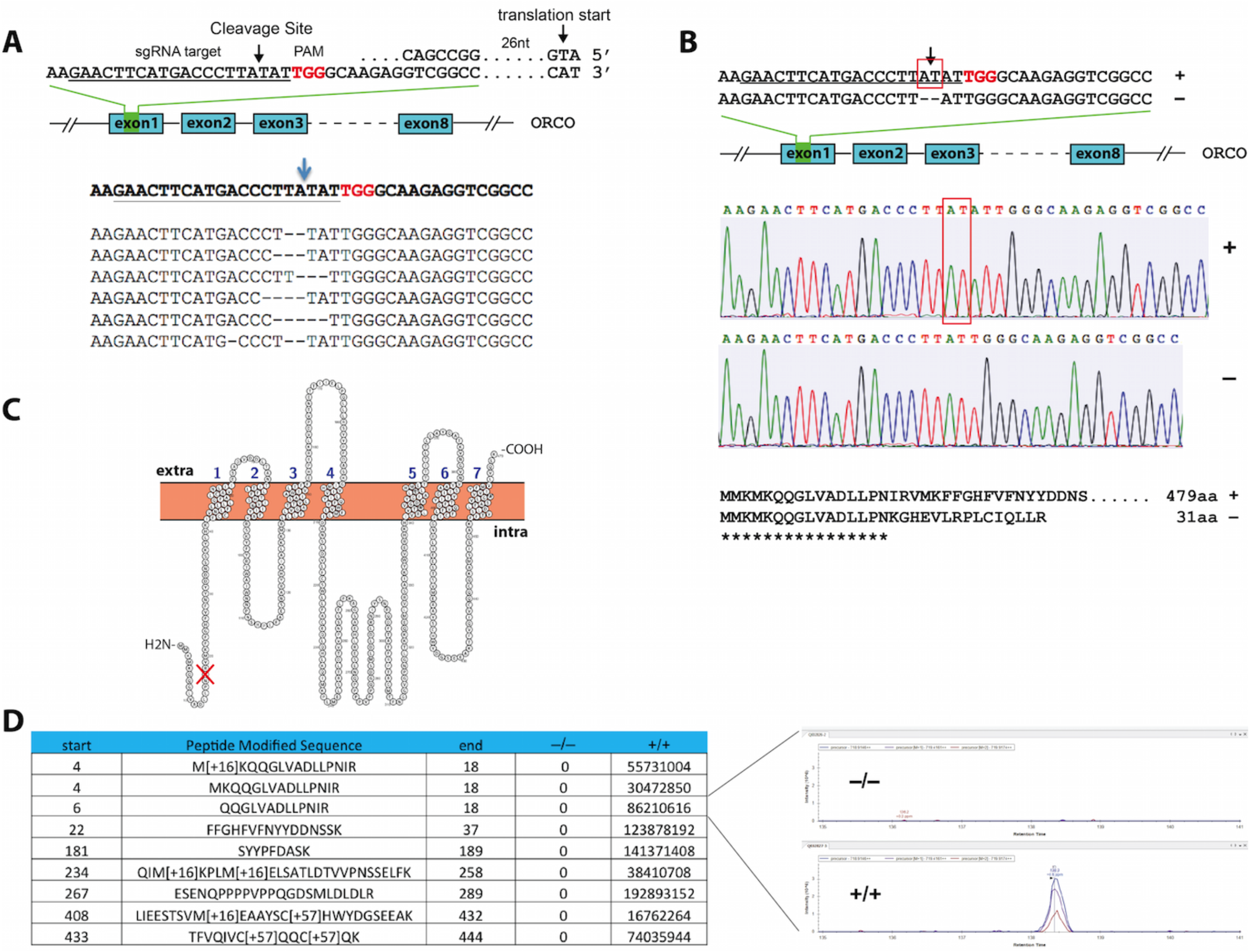
Development of somatic and germline mutations. (A) Microinjections of *orco* CRISPR into embryos generated multiple types of short deletions. The target site in the first exon of the *orco* gene, indicated by an arrow, is located 3 nucleotides upstream of the PAM (TGG) sequence. The small guide RNA (sgRNA) sequence is underlined. (B) The germline mutation used for this study was a 2-nucleotide AT deletion, as indicated in the sequencing chromatogram. The mutation caused a frameshift at codon 17 and premature stop codon, giving rise to the truncated protein of 31 amino acids. (C) The structure of Orco protein with 7 transmembrane domains, labeled 1 to 7. The mutation site is located at the N-terminus of the protein, indicated by the red mark. Extra and Intra: extra- and intra-cellular spaces. (D) Mass spectrometry indicates the abundance (peak area in inset) of detected peptides from Orco protein in the WT female ant compared to the F4 *orco* homozygous mutant ant. Each row represents a fragment of the WT Orco protein sequence with its start and end amino acid positions.

Among social insects, *Harpegnathos saltator* is uniquely suitable for the establishment and maintenance of mutant lines given its unusual caste plasticity (Yan et al., 2014). Specifically, any adult individual, most notably any adult mutant, can be converted to reproductive gamergate regardless of its caste at birth, and can therefore transmit the modified allele to its progeny. The gamergate phenotype can be induced in a social context by removing the queen (or the previously established gamergates) from a colony, or by completely isolating individual workers that, sensing the absence of the queen, become so-called solitary gamergates. Worker ovaries are atrophic and only the germarium can be observed (Fig. S1D), but grow to full size within 20 days after isolation when the solitary gamergate begins to lay eggs (Liebig et al., 1998; Peeters et al., 2000). Isolated workers are unmated and their unfertilized eggs develop into haploid males.

We converted isolated workers derived from injected embryos into gamergates (F0) and their haploid male progeny (F1) were genotyped by clipping their forewings and sequencing the CRISPR target site (Fig. 2B). One of the *orco* mutations comprised a deletion of two nucleotides in the first exon, resulting in a frame shift at codon 17 located at the N-terminal intracellular segment (Fig. 2C), and a premature stop codon (Fig. 2B), thus yielding a putative null mutant. The *orco* haploid mutant males did not display visible morphological or behavioral phenotypes and were fully fertile (see below). We back-crossed this animal to obtain F4 homozygous females (see below and Methods, Fig. 1C) and confirmed the absence of detectable Orco peptide in these homozygous individuals using mass spectrometry (Fig. 2D, Fig. S2C,D) and thus, that the frame shift resulted in a complete loss of function. In addition, the genetic inheritance followed Mendelian rules, as almost equal numbers of F3 wild type *vs*. mutant males, and F4 heterozygous *vs*. homozygous females were observed. Workers do not have wings and could be genotyped only after being sacrificed. Thus, all subsequent experiments were performed blind.

### Orco is required for chemical responses

In several insect species, *orco* mutants display a reduced response to general odorants (DeGennaro et al., 2013; Koutroumpa et al., 2016; Larsson et al., 2004; Li et al., 2016). To test the function of Orco in the response of workers to odorants, we established a bioassay that scores for antennal retraction after odorant delivery (Fig. S3A). In ants and other insects, seven odorants are known as alarm, or attractant/repellent pheromones (Boroczky et al., 2013; Capinera, 2004; Ditzen et al., 2008; Duffield et al., 1977; Zhao et al., 2010). These chemicals induce significant antennal retraction responses in WT *Harpegnathos* (Fig. S3B). 14 F4 putative *orco* mutant ants were paired (see Methods) with 14 WT control ants and their responses to different odorants were analyzed in parallel. While the WT ants responded robustly to these odorants, the F4 ants displayed bimodal responses, which allowed us to predict their genotypes. Statistical analyses for each odorant showed that the *orco* homozygous mutants displayed significantly lower responses to all odorants compared to heterozygous ants (Fig. 3A,B). Thus, genome editing of the *orco* locus resulted in the first loss-of-function mutation that was transmitted through the germline in an ant.

**Figure 3.**
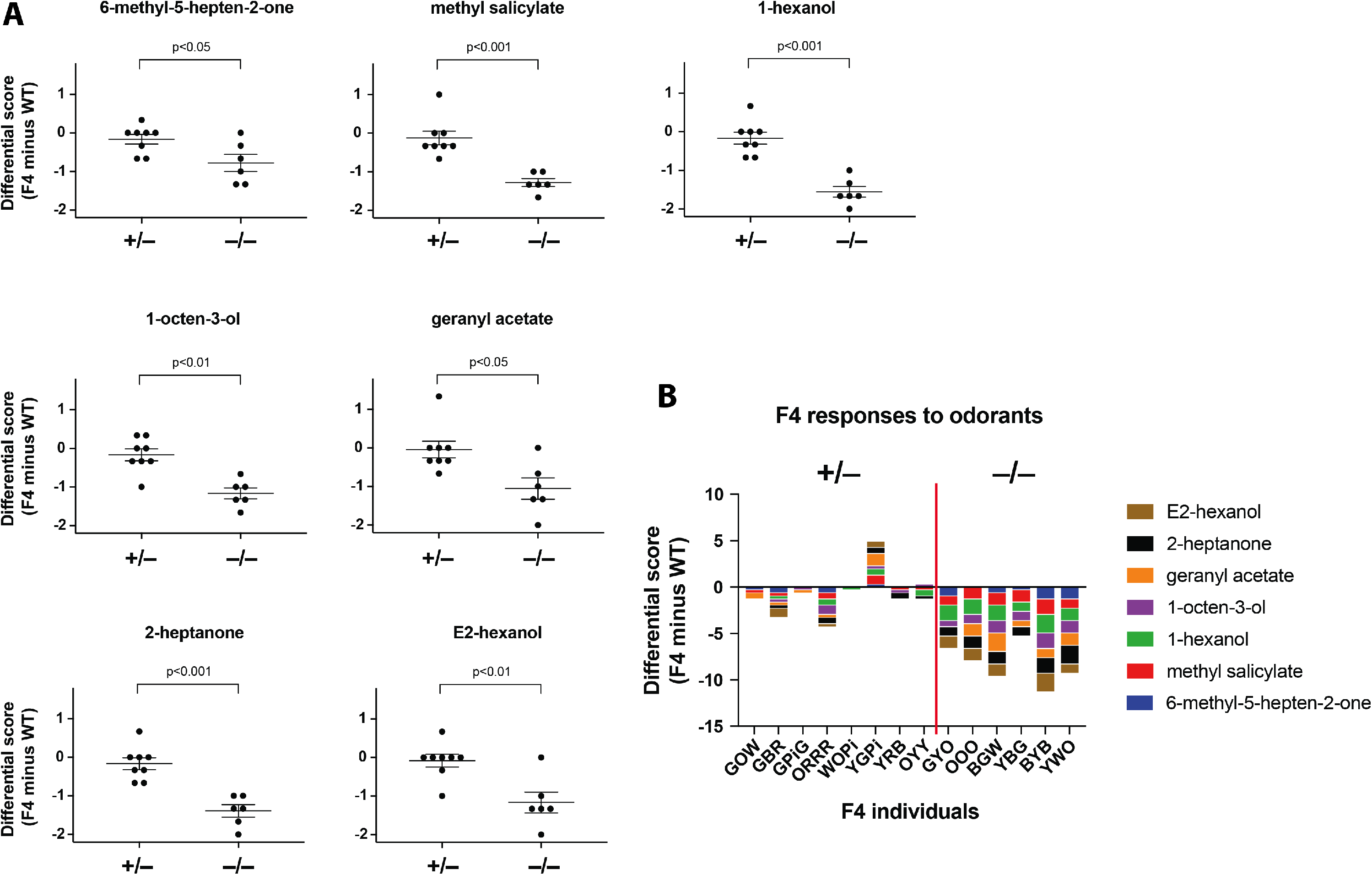
Reduced antennal responses to odorants in *orco* homozygous mutant ants. (A) The differential score (Y-axes) of each heterozygous (+/−, n=8) or homozygous ant (−/−, n=6) relative to its paired WT (+/+) ants was plotted and the p value indicated. Mann-Whitney test was used for statistical analyses. Error bars represent the standard error of the mean (SEM) in this figure and in other figures. (B) The cumulative differential scores in response to 7 odorants were separated into heterozygous and homozygous groups. Of note, YGPi was paired with an outlier, a WT ant with low responses to odorants.

### Orco mutant ants display behavioral phenotypes consistent with loss of pheromone sensing

Age-dependent polyphenism is a common phenomenon in eusocial insects. Young workers stay in the nest and nurse the brood, while older workers leave the nest and forage to bring food back to the colony (Haight, 2012; Robinson, 1987; Yan et al., 2014). *Harpegnathos* ants behave similarly since newly eclosed workers remain in the nest while older workers (>100 days old) leave the nest to forage. We speculated that chemical signals within the nest might act as “aggregation” pheromones for young workers (Ali and Morgan, 1990; Depickere et al., 2004). Interestingly, young homozygous *orco* worker ants (age <50 days) spent a much longer time outside the nest than WT and heterozygous young workers (Fig. 4A), suggesting that the *orco* mutation caused a measurable phenotype in a distinct social behavior (leaving the colony to forage), most likely due to a defect in sensing chemical signals in the nest. This wandering phenotype was not due to an early transition to foraging in homozygous *orco* mutants since the ants that spent much of their time outside the nest never foraged to return crickets back to the nest (Fig. 4B). In fact, these ants fed on crickets brought to the nest by WT workers. This discrepancy suggests that *orco* mutants are not only impaired in their ability to detect “aggregation” pheromones, but also in sensing the presence of prey, such as crickets, when outside of the nest.

**Figure 4.**
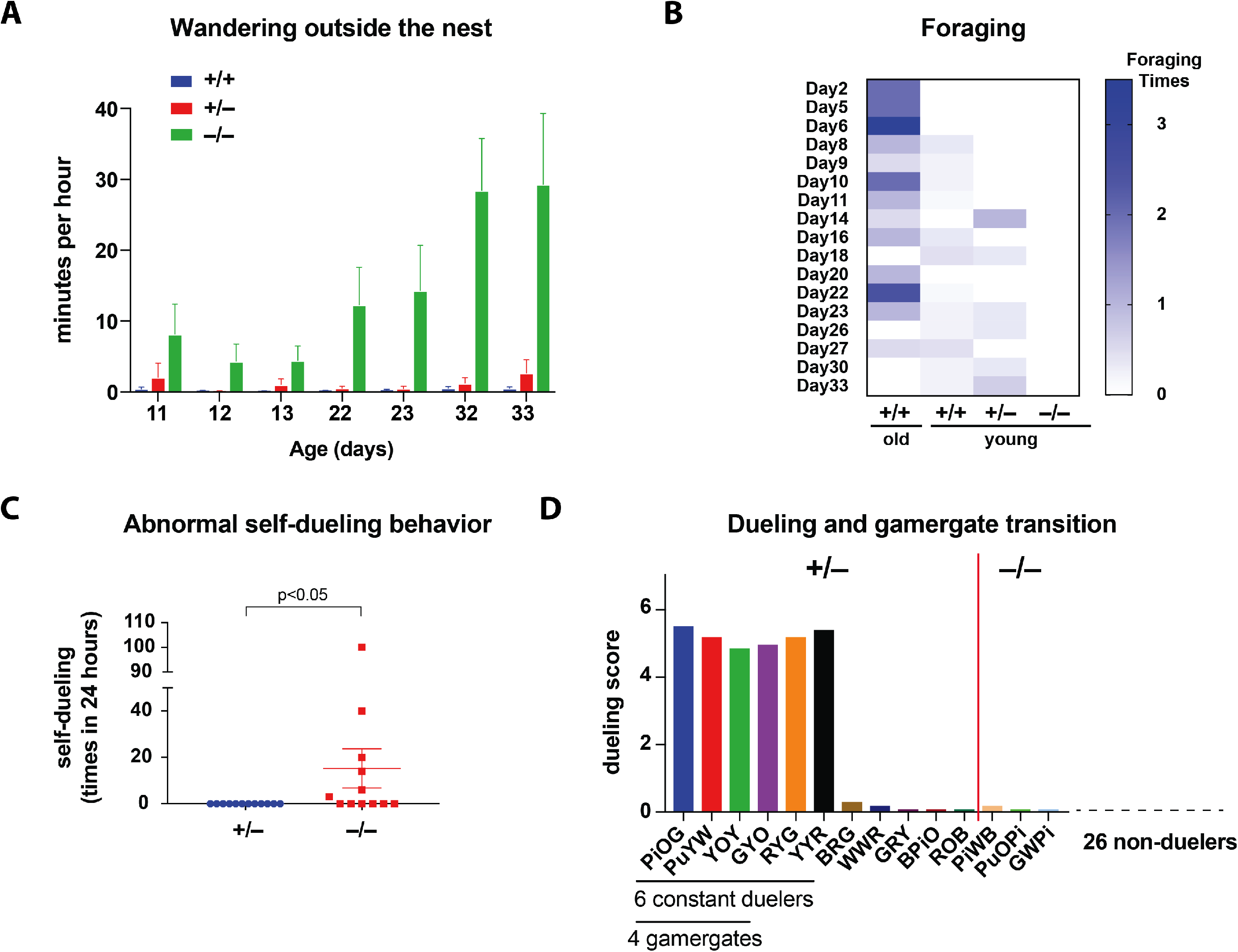
Behavioral phenotypes in *orco* homozygous mutant ants. (A) The time for homozygous young workers (<50 days old, n=6) vs. heterozygous (n=6) and WT young workers to wander outside of the nest: p<0.001 between homozygotes and heterozygotes; p<0.0001 between homozygotes and WTs; p=0.9335 between heterozygotes and WTs (Two-way ANOVA with Tukey’s multiple comparisons test). (B) The wandering homozygous ants did not forage. The foraging times are indicated in the heat map: p<0.05 between young homozygotes and old WT workers (Two-way ANOVA with Tukey’s multiple comparisons test). (C) Abnormal self-dueling behavior was found in 6 out of 12 individually isolated homozygotes, but not in any of the 12 heterozygotes, under the same conditions: p<0.05 (Mann-Whitney test). (D) The average dueling times in 11 heterozygous duelers (left) vs. 3 homozygous duelers (right) in the first 9 dueling days. The 6 constant duelers and 4 gamergates are indicated under the individual ants.

### Loss of orco impairs worker dueling and transition to gamergates

In the absence of a reproductive queen or gamergates, a large proportion of workers (30-60%) enter a ritualistic dueling behavior. While most desist rapidly, those that remain constant duelers finally become gamergates and lay eggs (Sasaki et al., 2016)(G.M., in preparation)(J. Gospocic in preparation). To address whether the *orco* mutation affected dueling and the gamergate transition, we generated a colony with 39 randomly selected young F4 putative *orco* mutant ants within a setting devoid of a queen or gamergates, and tested for their ability to become gamergates. As usual, dueling started 3 days after colony setup. During the first day of dueling, 14 out of 39 workers (36%) dueled. This number became lower with time as some duelers returned to worker behavior. From dueling day 3 to day 9, only 6 workers constantly dueled. Finally, 4 of them became gamergates (Fig. 4D). Subsequent genotype analyses indicated that only 3 out of the 17 homozygous *orco* mutants (18%) entered dueling, but they quickly ceased on day 2 and never became gamergates (Fig.4D). In contrast, 11 out of the 22 heterozygotes (50%) joined dueling. All 6 remaining duelers and the 4 final gamergates were all heterozygous. This suggests that loss of Orco resulted in the inability to properly sense the social environment and participate in appropriate dueling behavior to achieve reproductive status (Liebig et al., 2000).

In a solitary environment, the homozygous *orco* mutants also displayed another abnormal behavior: they quickly moved their antennae, similar to the quick antennal movement observed during dueling with other workers in the transition to gamergate. This “self-dueling” behavior was never observed in isolated heterozygotes (Fig. 4C). This aberrant behavior is paradoxical given that the homozygous mutants rarely engaged in dueling with heterozygous workers when in a society (see above), underscoring that a functional Orco protein is required to correctly sense and respond to environmental cues (see above).

### Mating and reproduction are impaired in homozygous orco mutants

In *Harpegnathos*, the mating ritual begins with the female grooming the male for a few hours, until copulation begins and lasts for 20-50 seconds. *Harpegnathos* males can mate multiple times (polygyny) but females mate only once (monoandry). Heterozygous females, like WT, groomed and mated with males. However, we never observed any grooming and mating behavior from homozygous mutant females, consistent with the hypothesis that sex pheromones used for attracting mates are not sensed by *orco* mutant females.

When individually isolated, the *orco* homozygous ants, like heterozygous, were capable of laying haploid eggs, but they displayed reduced fecundity. They started to lay eggs significantly later than heterozygotes (median: 39 days in homozygotes *vs*. 29.5 days in heterozygotes) (Fig. 5A) and laid 7 times fewer eggs during the first 50 days after emergence (Fig. 5B). Moreover, the homozygotes did not properly care for their brood. Although one *orco* homozygote produced larvae (Fig. 5C), these larvae failed to survive and develop to pupae. In contrast, all isolated heterozygous gamergates produced larvae (Fig. 5C), pupae and adults. Interestingly, *orco* mutant males, like WT males, were capable of mating with females to have healthy progeny, which allowed us to propagate the mutants. This suggests that, unlike in females, Orco is not required in males for mating and proper reproduction, consistent with the much lower number of *Or* genes expressed in male sensilla (McKenzie et al., 2016; Zhou et al., 2012) and thus, a reduced importance for OR-mediated chemosensation in contributing to the more defined male behaviors.

**Figure 5.**
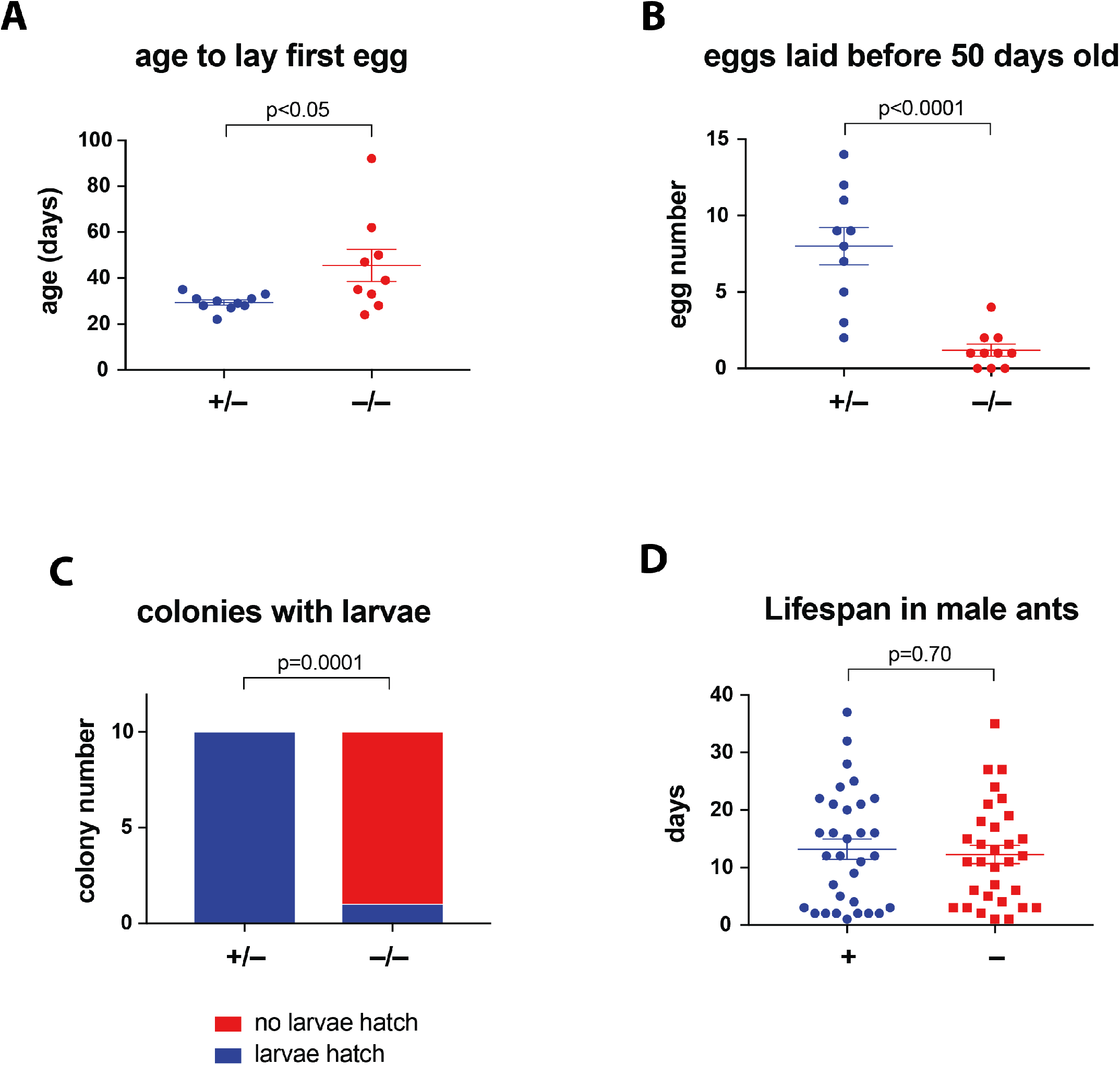
Defective reproduction and nursing in *orco* homozygous mutant females, but no change in lifespan of *orco* mutant males. (A) When individually isolated, the heterozygous ants (n=10) laid the first egg at day 29.4 ± 1.1, while the homozygous mutant ants (n=9) did so at day 45.6 ± 7.1: p<0.05 (unpaired t test). (B) The heterozygous ants (n=10) laid 8.0 ± 1.2 eggs during the first 50 days after eclosion, while the homozygous ants (n=10) laid 1.2 ± 0.4 eggs before 50 days old: p<0.0001 (unpaired t test). (C) While all heterozygous ants (n=10) produced larvae, only 1 homozygous ants (n=10) did so: p=0.0001 (Fisher’s exact test). (D) The WT (+, n=32) and hemizygous mutant F3 male ants (−, n=31), which lived with individually isolated female ants, had average lifespans of 13.19 ± 1.76 days and 12.26 ± 1.58 days, respectively: p>0.05 (unpaired t test).

Homozygous *orco* mutant females were viable and healthy, but we could not evaluate their lifespan since most F4 individuals were heavily manipulated for behavioral experiments and because of the long natural lifespan of *Harpegnathos* females (median lifespan: ~7 months)(Haight, 2012). However, we did measure the lifespan of male mutants, finding it similar to that of WT males that exhibit a shorter lifespan than females (Fig. 5D), suggesting that impaired OR function does not reduce survival of male ants, and that no significant off-target mutations affect their viability.

### Mutation in orco leads to dramatic decreases in number and in size of antennal lobe glomeruli

In *Drosophila* and in mammals, all ORNs expressing the same *Or* gene project to the same specific glomeruli in the antennal lobes or olfactory bulb, respectively (Mombaerts, 2006; Vosshall et al., 2000). Each *Drosophila* ORN is believed to express only one *Or* genes and projects to one of 42 distinct glomeruli. Mutations in *orco* or manipulation of ORN activity do not alter the morphology of the AL where ORNs project (Chiang et al., 2009; Larsson et al., 2004). This led to the assumption that, unlike in mammals, ORN activity is not required to establish and maintain the number and shape of glomeruli that appear to be hardwired. We were therefore surprised that, while performing our studies, a similar *orco* mutant in another species of ants, *Ooceraea biroi*, a parthenogenetic ant with a peculiar phase-alternating foraging and egg laying behavior, was described in bioRxiv (CSHL) as exhibiting a dramatic decrease in the number of glomeruli (Trible et al., 2017) (Daniel Kronauer, personal communication).

We dissected the brain of F4 ants and performed staining using Phalloidin to mark the glomeruli (Hoyer et al., 2005). The volume of the AL of *orco* homozygous *Harpegnathos* mutants (2.60 ± 0.18 × 10^6^ μm^3^) was much smaller than that of heterozygous (4.29 ± 0.16 × 10^6^ μm^3^) or WT ants (4.26 ± 0.24 × 10^6^ μm^3^) (Fig. 6A,B, Fig. S4A). Moreover, as was observed in *Ooceraea orco* mutant ants (Trible et al., 2017), *Harpegnathos orco* mutants had a dramatically reduced number of glomeruli: 275 glomeruli were present in WT and 270 in heterozygous, while only 62 were observed in homozygous *orco* mutants (Fig. 6C). Strikingly, in homozygous *orco Harpegnathos* mutants, the diameter of the remaining individual glomeruli was 1.6 times that of heterozygous (Fig. 6D). The dramatic morphological changes in glomeruli and antennal lobes in *orco* mutant ants suggest that these ants, in contrast to *Drosophila*, display strong olfaction-mediated neuroanatomical plasticity. The increased size of the glomeruli in the mutant suggests that individual glomeruli remain merged in the absence of activity and that ORNs innervate a small number of remaining glomeruli that might develop, as in *Drosophila*, in the absence of activity.

**Figure 6.**
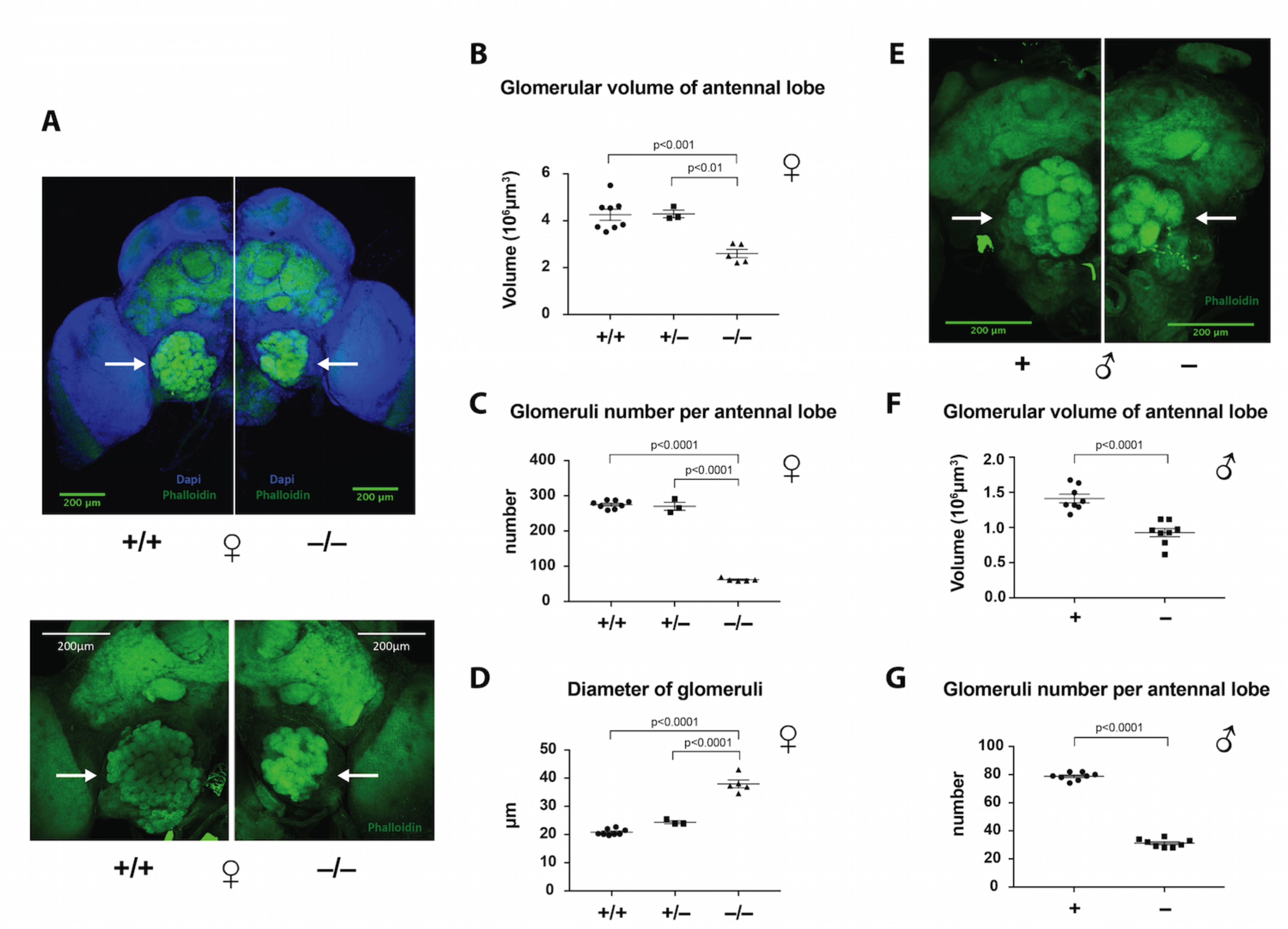
Neuroanatomical phenotypes in female and male mutant ants. The ant brain samples were stained with Phalloidin (green) and counter-stained with DAPI (blue). Confocal images of two WT vs. two homozygous mutant ants are shown in (A): 10X objective (top panel) and 20X objective (bottom panel). Confocal images of WT vs. mutant male ants using 20X objective are shown in (E). The glomerular areas of antennal lobes are indicated by arrows in (A) and (E). (B) The average glomerular volume of left and right antennal lobes in WT (n=8): 4.26 ± 0.24 × 10^6^ μm^3^, heterozygous (n=3): 4.29 ± 0.16 × 10^6^ μm^3^, and homozygous female ants (n=5): 2.60 ± 0.18 × 10^6^ μm^3^. (C) The number of glomeruli in WT: 275.2 ± 4.0, heterozygous: 270.2 ± 11.2, and homozygous female ants: 61.8 ± 1.9. (D) The diameter of glomeruli in WT: 20.8 ± 0.4 μm, heterozygous: 24.4 ± 0.6 μm, and homozygous female ants: 38.0 ± 1.4 μm. (F) The glomerular volume of each antennal lobe in WT (n=8): 1.41 ± 0.06 × 10^6^ μm^3^, and mutant male ants (n=8): 0.93 ± 0.06 × 10^6^ μm^3^. (G) The number of glomeruli in WT: 78.8 ± 1.0 μm, and mutant male ants: 31.3 ± 1.0 μm. Female: one-way ANOVA plus Tukey’s multiple comparisons test; male: unpaired t test.

The size of the AL of *orco* hemizygous mutant males was also significantly smaller (0.93 ± 0.06 × 10^6^ μm^3^) than in WT males (1.41 ± 0.06 X 10^6^ μm^3^) (Fig. 6E,F). The number of glomeruli was also dramatically decreased: 31 vs. 79 glomeruli in WT (Hoyer et al., 2005) (Fig. 6G). Male glomeruli widely varied in size with dorsal glomeruli being much larger than ventral ones (Hoyer et al., 2005). In *orco* mutant males, the dorsal glomeruli became smaller (Fig. S4B,C), while some ventral glomeruli did not display a significant change in size (Fig. S4C). Since *orco* mutant males exhibit normal mating, fertility and lifespan, the remaining glomeruli are apparently sufficient for their reproductive behavior.

## Discussion

Reproductive division of labor, the defining feature of eusocial animals, poses a significant challenge to generating and maintaining germline mutations in laboratory colonies because only a few individuals in a colony reproduce. The advantage of *Harpegnathos* over many other hymenopteran eusocial species is that all females can sexually reproduce after conversion to gamergate, which can be induced by social isolation (Yan et al., 2014). This characteristic provides the opportunity to effectively use gene targeting to study developmental, neuronal and behavioral plasticity in a social insect. In parallel to our studies, Daniel Kronauer and his colleagues (Trible et al., 2017) developed lines of *orco* mutants using thelytokous parthenogenesis in *Ooceraea* where all progeny are maternal clones. The phenotypes they obtained with this very different species that diverged from *Harpegnathos* 120 millions years ago (McKenzie et al., 2014), are fully consistent with our own observations. Thus, taken together, these two studies describe the first genetic mutants in eusocial insects. *Harpegnathos*, in particular, provides an apt species to utilize as a genetic model organism as the ability to perform genetic crosses will foster the development of more sophisticated genetic tools similar to those available in *Drosophila* (e.g. UAS-GAL4 inducible lines, multiple mutants, etc.). Such a repertoire would then allow for an expansion of genetic analyses to further our understanding of the genetic and epigenetic regulation of social behavior.

In ants, queen pheromones play a major role in suppressing worker reproduction (Van Oystaeyen et al., 2014). During the worker to gamergate transition in *Harpegnathos*, the CHC profile shifts to longer chain hydrocarbons, which mimic the CHC profiles of the queen. Some of these long chain hydrocarbons most likely act as queen pheromones (Liebig et al., 2000), the absence of which triggers dueling in workers. The *orco* mutant workers display a series of behavioral phenotypes that are consistent with the loss of olfactory function: 1) they wander out of the social group and are unable to forage successfully; 2) they appear to be largely unable to communicate with conspecifics, a feature that is reminiscent of Autistic Spectrum Disorder; 3) they exhibit a dysfunctional behavior in isolation, consisting of quick movement of antennae that resemble dueling despite the absence of other ants; 4) they fail to fully engage in dueling and never transition to gamergates in the presence of other ants; 5) they do not produce progeny because they lay very few eggs and do not tend to them; 6) they ignore the presence of males and fail to mate, presumably because they cannot detect the male sex pheromones.

In addition, and similar to the observations in *Ooceraea* (Trible et al., 2017), loss of Orco-dependent OR signaling resulted in a dramatic decrease in the size of the antennal lobes in both males and female *Harpegnathos*, as well as in the number of glomeruli, linking neuroanatomy to Orco function in these insects. Only a small proportion (31 out of 79 in *Harpegnathos* males, 62 out of 275 in females) of glomeruli remained in the absence of *orco* function. Interestingly, the number of remaining glomeruli is similar to that (42) in *Drosophila*, which is hardwired and Orco-independent. This raises the interesting possibility that ants pattern their antennal lobe through two distinct mechanisms: one that is Orco-dependent and another that is ancestral in insects and is independent of Orco function, as in *Drosophila*. The latter might correspond to the remaining glomeruli in our mutants. This type of patterning is possible with the limited number of ORNs that can connect to their cognate glomeruli using typical guidance molecules such as protocadherins or Dpr/DIPs. However, with the amplification of the 9-exon OR family in ants (McKenzie et al., 2016; Zhou et al., 2012), it would not be possible to pattern several hundred ORNs with a Sperry type of mechanism (Sperry, 1963); the activity of the ORNs is thus required as a second mechanism that might split proto-glomeruli into multiple glomeruli as a function of such activity. The presence of considerably fewer glomeruli in males correlates with the lack of expression of 9-exon ORs and thus, reflects less reliance on activity-dependent patterning.

A similar mechanism has been proposed for the formation of the more than 1,000 pairs of glomeruli in mice that require the activity of ORs. The mechanisms that mediate the projection of ORNs to specific glomeruli remain unknown. It should be noted that the structure of ORs in insects, which are cation channels, is very different from that of mammalian ORs, which are GPCRs. Yet, the two signal transduction cascades might converge to direct the neurons to their target glomeruli. In mammals, although the gross anatomy of the adult brain does not change dramatically in the absence of electrophysiological activity, significant changes can be observed in several parts of the brain when activity is prevented during critical periods (Espinosa and Stryker, 2012; Hubel and Wiesel, 1979). Furthermore, neural stem cells (NSCs) that contribute to adult neurogenesis in two brain regions, the hippocampus and the olfactory bulb (OB), are regulated by environmental input throughout life (Suh et al., 2009). In contrast, the insect brain is widely believed to be hardwired and to depend only minimally on activity patterning (Hassan and Hiesinger, 2015; Jefferis et al., 2001). Therefore, the *orco* phenotype in ants might be one of the first examples of activity-driven patterning in insects (Petsakou et al., 2015). In other insects, such as the moth *Manduca sexta*, odorant-induced ORN activity appears after the development of the 64 glomeruli (Oland and Tolbert, 1996) and thus, is unlikely to play a role.

Future investigations into *Harpegnathos orco* mutants will address the development of the ORNs, and at what stage during development or after eclosion is ORN activity required for the establishment and maintenance of glomeruli. To further assess whether activity-dependent neuroanatomical plasticity is conserved in insects, it will be useful to analyze non-eusocial insects, like the solitary parasitoid wasp *Nasonia vitripennis*, which also possesses a much larger number of *Or* genes than *Drosophila* (Zhou et al., 2012). Of note, CRISPR-mediated mutagenesis in *Nasonia* has been achieved in our laboratory (M. Perry and C.D. personal communication).

An alternative to the model that ants exhibit two modes of patterning of ORNs is that the remaining glomeruli in the *orco* mutant ants are those innervated by GR (gustatory receptor)-or IR (ionotropic receptor)-expressing ORNs, which remain functional in the absence of *orco*. In mosquitos and *Drosophila*, a significant proportion of glomeruli are targeted by neurons expressing GRs or IRs (Benton et al., 2009; Jones et al., 2007; Riabinina et al., 2016). In fact, significant chemosensory responses still remain in the *Harpegnathos orco* homozygous workers, as these mutants, like their heterozygous and WT counterparts, showed timid behavior towards the reproductive gamergates, and did not duel or lay eggs when in a stable colony.

The behavioral flexibility of *Harpegnathos* ants presents unique experimental opportunities to dissect the molecular mechanisms by which social and neuronal plasticity are regulated (Bonasio, 2012; Bonasio et al., 2010; Yan et al., 2014). The findings in the *orco* mutants illustrate how the olfactory system and chemosensation play critical roles in ant sociality. The availability of a high quality genome sequence (Bonasio et al., 2010), DNA methylome (Bonasio et al., 2012), and now genome editing tools effectively raise the status of *Harpegnathos* to that of a model organism uniquely suited for investigating the genetic and epigenetic regulation of behavior as well as the regulation of neural plasticity.

## Materials and Methods

### Regular maintenance of Harpegnathos ant colonies

*Harpegnathos saltator* colonies were initially transported from Jürgen Liebig’s laboratory at Arizona State University. Ants were maintained in plastic boxes (Pioneer Plastics, Inc.) inside the USDA approved ant room at 22-25°C. Small boxes (9.5 × 9.5 cm^2^) were used to rear individually isolated workers. Medium (19 × 13.5 cm^2^) and large boxes (27 × 19 cm^2^) were used to rear social colonies. The floor of the boxes was made of plaster, and a glass was used in each medium or large box to separate the inside of the nest where the reproductive females and young workers (nurses) lived with the brood from outside of the nest that was the foraging area for old workers (foragers). Colonies were fed with live or pre-stung crickets 3-6 times a week. Crickets were pre-stung by workers in the large colonies to provide to the individually isolated ants.

### Embryos injections and post-injection maintenance

Quartz glass needles were made using the P2000 Micropipette Puller (Shutter Instrument). Embryos were lined on the double-sided tape and microinjected with Cas9 proteins (0.2 μg/μl) and *in vitro* synthesized small guide RNAs (sgRNAs, 0.2 μg/μl) using the Eppendorf FemtoJet II microinjector. The design of the sgRNAs was described previously (Perry et al., 2016). The *orco* gene sequence (Hsal_00588) was obtained from the BGI DNA database and the genome sequence of *Harpegnathos saltator* was reported (Bonasio et al., 2010). The structure of transmembrane domains in the Orco protein was analyzed by Protter (Omasits et al., 2014).

We first found that most injected embryos were destroyed by workers after they were reintroduced to the colony. To increase the survival rate, we optimized the method in rearing the injected embryos on 1% agar plates with 2% Antibiotic-Antimycotic (ThermoFisher Scientific) after 70% ethanol treatment. After one month, when the first embryo hatched, they were transported to a nest with a few helper workers, as worker nursing is essential for the survival of larvae. The remaining embryos hatched in the colony. We fed them with pre-stung crickets to maintain stability in the small nest with injected embryos.

### Genotyping of embryo, larval and adult tissues

Different methods were used to analyze the genotypes of different stages of ants. First, to test somatic mutation, embryos were harvested 3 days after injections and 5 embryos were pooled for genomic DNA extraction and PCR amplification using the Taq DNA polymerase. PCR products were cloned to pGEM vector with A-T overhang (Promega). Several clones were sequenced. Second, amplified libraries were generated from extracted genomic DNA followed by MiSeq at the NYU Genomic Center to analyze the efficiency. Third, to test the germline transformation, single F1 larvae were harvested for DNA sequencing. Fourth, to test the genotype in adult males, their forewings were cleaved for genomic DNA extraction and sequencing, while the individuals were alive and capable of mating with females. Fifth, to test the genotype in adult females, each individual was harvested and part of its thorax tissue was genotyped. PCR primers for genotyping:

Forward primer: TCCCGTATCGGCGATAACGATGACAAGC

Reverse primer: TGACCGATAGACTAAGACAGGCCG

### Maintenance of the mutant allele and development of homozygous mutant ants

In most eusocial insects, mating cannot be achieved in the laboratory (Yan et al., 2014), but it can be carried out in *Harpegnathos* (Liebig et al., 1998). Indeed, we have successfully recorded the mating of a single male with a single virgin female of *Harpegnathos* in the lab. This controlled mating allowed us to generate homozygous mutants in the F4 generation.

The mutant F1 males were identified by genotyping. Each mutant male was paired with a WT virgin female for mating under the video camera. When the mating was confirmed, the male was transferred to another nest with a new WT female to maximize the number of progeny. The male could inseminate multiple females from day 0 to day 7 after emergence. No mating was observed after day 7. The inseminated WT females gave rise to F2 heterozygotes, which in turn generated F3 hemizygous males. The mutant F3 males were backcrossed to F2s to generate F4 homozygotes (Fig. 1C). <u>Of note, as the F4s were either heterozygotes or homozygotes, the assays were performed blind without knowing their genotypes until they were harvested for genotyping.</u>

### Mass spectrometry

Proteins were extracted from antennae cleaved from WT and homozygous mutant ants. The protein extract was run on SDS-PAGE gels. Gel slices were trypsin digested followed by liquid chromatography-tandem mass spectrometry (LC-MS/MS) analysis performed at the Biological Mass Spectrometry Facility at Rutgers, The State University of New Jersey. Skyline software was used for the quantification of peak area.

### Immunohistochemistry of embryo nuclei and adult brains

Embryos were harvested every 4 hours from 12 hours after egg deposition (AED) to 40 hours AED. After harvesting, the embryos were boiled for 45 sec, quenched quickly on ice, and fixed with 4% paraformaldehyde (PFA) supplemented with DMSO and heptane (Khila and Abouheif, 2009), followed by protocols for *Drosophila* embryo sectioning in the lab.

Ant brain tissues were dissected and fixed in 4% PFA and washed three times using 1X PBS with 0.3% Triton X-100. The tissues were incubated with DAPI (2 μg/μl) and Alexa Fluor^®^ 488 Phalloidin (1/80, ThermoFisher Scientific) at room temperature for 2 hours, and then mounted and scanned using Confocal Leica TCS SP5 microscope at NYU. Images of AL glomeruli were generated using Z project in Fiji (or ImageJ).

### Behavioral responses to multiple chemicals

Each F4 ant was painted with multiple color dots on their thoraces and thus named by the respective color combination (*e.g*. GOW was 3 colors of Green, Orange and White from anterior to posterior). Each pair of WT and F4 ants was immobilized on their thoraces (Fig. S3A). Their antennae were able to move without restrictions. 20μl of a 1% odorant/paraffin solution was transferred onto a small piece of filter paper inside a glass pipette. The odorants were delivered as a pulse into a constant 1.7l/min flow of air supplied by the building pressure gas system. The one second pulses with a 280ml/min flow rate were controlled using a Syntech Stimulus Controller, model CS-55. The air flew directly into a Y-shaped glass tube with an inner diameter of approximately 15mm. An F4 was placed in one arm of the Y-tube and a wild type into the other. The control stimuli (solvent alone) were delivered before the chemical stimuli. The antennal responses to odorant and control stimuli were video-recorded and each ant was tested for 3 rounds of the complete odorant panel. The sequence of odorants was randomized for each test, as was the position of the WT and F4 ants in the Y-tube. The videos were analyzed blind (without knowledge of the ant genotypes) and scored as follows: 0 for no retraction (or random antennae movement), 1 for mild retraction (slow or incomplete retraction), and 2 for strong retraction (quick and complete retraction). Each differential score was calculated using the score of F4 ant minus the score of paired WT ant. The Mann-Whitney test was performed for statistical analyses using GraphPad Prism 7 (version 7.00 for MacOS, GraphPad Software, La Jolla California USA). To predict the genotype, their responses to seven odorants were added up: 7 homozygous vs. 7 heterozygous ants (1:1 ratio) were predicted based on their differential scores.

### Analyses of ant behavior, egg laying and lifespan

The nests for *Harpegnathos* were separated into two areas, inside the colony (under glass) and outside of the colony. F4 ants were grouped with WT ants that included gamergates, young workers (nurses) and old workers (foragers). Gamergates constantly stayed inside with young workers and brood, while old workers brought pre-stung crickets from outside to inside. The nests were videorecorded and the time for each worker in staying outside of the colony was counted. Two-way ANOVA plus Tukey’s multiple comparisons test were performed for statistical analysis.

To observe behaviors, reproduction and lifespan for solitary female ants, individual F4s were isolated to generate gamergates. Their behaviors were video-recorded and their brood was counted weekly. The videos were analyzed manually and blind. Mann-Whitney and Fisher tests were performed for statistical analyses of abnormal antennal movement, individual egg-laying and larvae-containing colonies. For the lifespan analysis in males, 32 WT and 31 hemizygous mutant ants in the F3 generation were compared, and the unpaired t test was performed for statistical analysis.

To address the role of Orco in the dueling and gamergate transition, we generated a colony (n=39) with homozygous and heterozygous young workers, without gamergates. The frequency of dueling was observed 6 times per day (each time for 30 min) during the first 9 days after dueling started. The gamergates were determined by their egg-laying 3 months after separation. The statistical analysis of dueling and gamergate transition in the heterozygous vs. homozygous ants was performed using Fisher’s exact test, and the dueling frequencies in the heterozygous and homozygous duelers was performed using Wilcoxon matched-pairs signed rank test.

### Measurement of antennal lobes

Confocal microscopy was used to visualize whole mount *Harpegnathos* brains using DAPI and Alexa Fluor^®^ 488 Phalloidin. Glomeruli were reconstructed from image stacks using the segmentation editor of the Fiji (Schindelin et al., 2012). The AL areas were manually determined by the outlines of the cluster of glomeruli. The glomerular volume of each AL was analyzed by the Object Counter 3D Fiji plugin (Bolte and Cordelieres, 2006), and normalized by the volume of central region (protocerebrum) of the same brain, including protocerebral neutrophil, vertical lobe, peduncle and central bodies (Hoyer et al., 2005). The volume of the left and right ALs was averaged to find each individual’s AL volume. For glomeruli counting, each glomerulus was identified in the image stack and tracked with a manually placed high intensity mark using Fiji. The glomeruli counts in the left and right ALs were averaged to obtain the count per lobe in each organism. To measure the size of glomeruli, an AL per individual ant was reconstructed from image stacks using Fiji. Multiple glomeruli in each AL were picked at random (27 glomeruli per female ant, and 3-14 per male ant depending on the location of glomeruli). The 3D image was calibrated with zoom settings and a line was drawn across the shortest midsections. The length of each line was measured using Fiji. Similar methods were used to quantify the glomerular volume and number of glomeruli in eight ALs in male ants. One-way ANOVA plus Tukey’s multiple comparisons test were performed for female ants and unpaired t test for male ants in the statistical analyses using GraphPad Prism 7.

## Acknowledgements

We appreciate the advice from Gregory Pask and Daniel Simola in designing *orco* sgRNAs and improvement of reproduction in individually isolated ants, from Brittany Enzmann on embryo injections, from Waring (Buck) Trible (Daniel Kronauer’s lab) on the deep sequencing to test somatic mutation efficiency, from Haiyan Zheng and Shengjiang Tu on mass spectrometry, and from Lynne Vales on manuscript preparation. We also appreciate the assistance from Heike Pelka, Anika Paradkar, Nicholas Rogers and Dong Chen in managing the ant room, maintaining ant colonies or analyzing behavioral videos.

This work was supported by a Howard Hughes Medical Institute Collaborative Innovation Award (HCIA) #2009005 and NIH R21 Exploratory/Developmental Research Grant R21GM114457. H.Y. was a NIH Ruth L. Kirschstein NRSA Postdoctoral Fellow (F32AG044971). R.B. was supported by the Searle Scholars Program and by an NIH New Innovator Award (DP2MH107055).

## Supplemental Figure Legends

**Figure S1.**
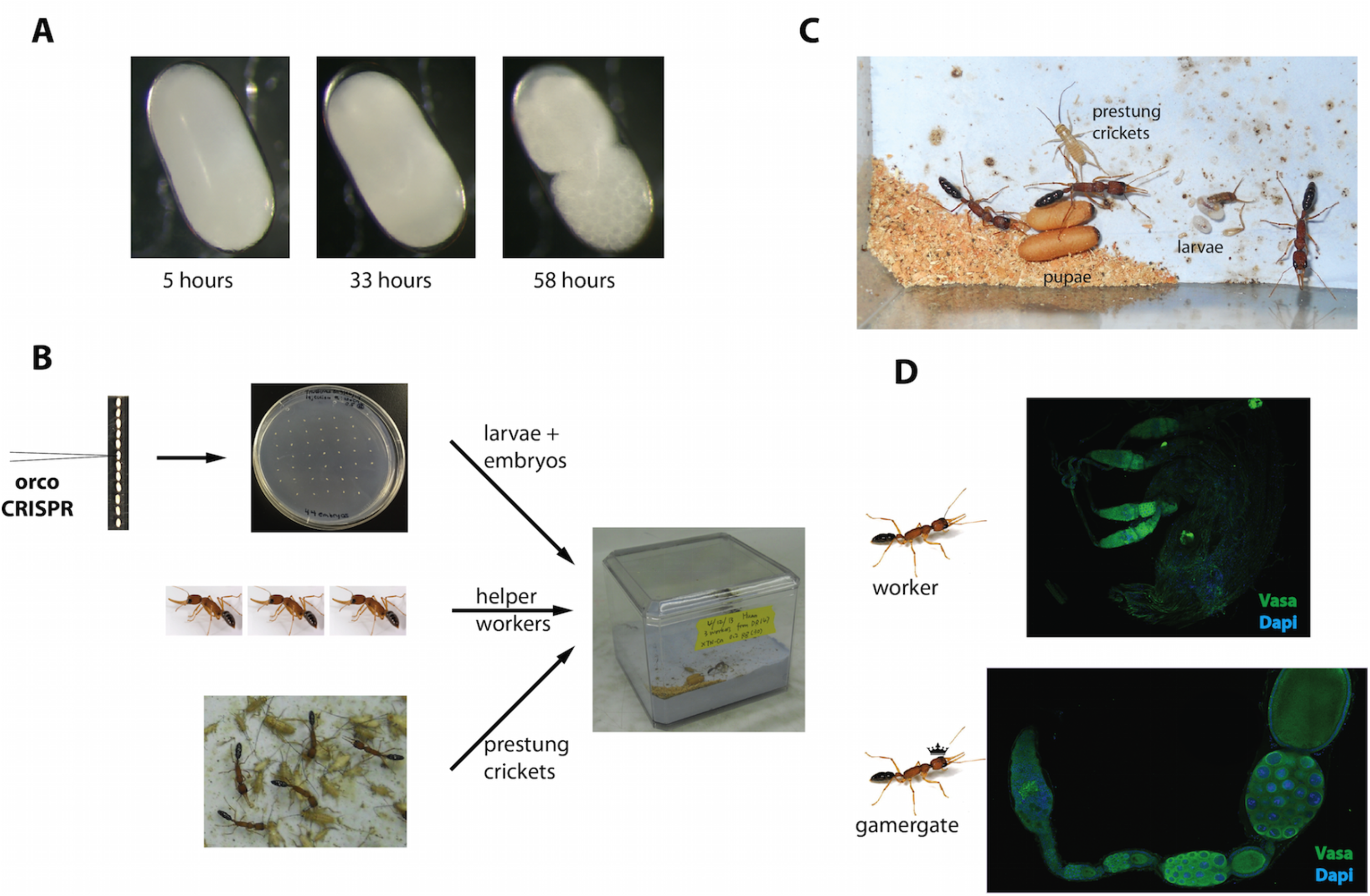
Embryo development, injections, maintenance and the worker-to-gamergate transition in *Harpegnathos*. (A) A *Harpegnathos* young embryo was placed in the mineral oil. Photos were taken under a dissection microscope at different time points after egg deposition (AED). The contraction of vitelline membrane from the chorion at 33 hours AED indicated that the embryo was still at the syncytial stage (Wilt and Wessells, 1967). The embryo at 58 hours AED reached gastrulation. (B) After injections, embryos were removed from the double-sided tape and placed on an agar plate. After one month, they were transported to a colony with 3 nursing workers. The colony was fed with pre-stung crickets. (C) Later on, the larvae and pupae were developed from the injected embryos. (D) The ovaries dissected from worker vs. gamergate indicated that egg chambers and oocytes grow in the gamergate, while the ovary growth is arrested in the worker. Vasa: green; DAPI: blue.

**Figure S2.**
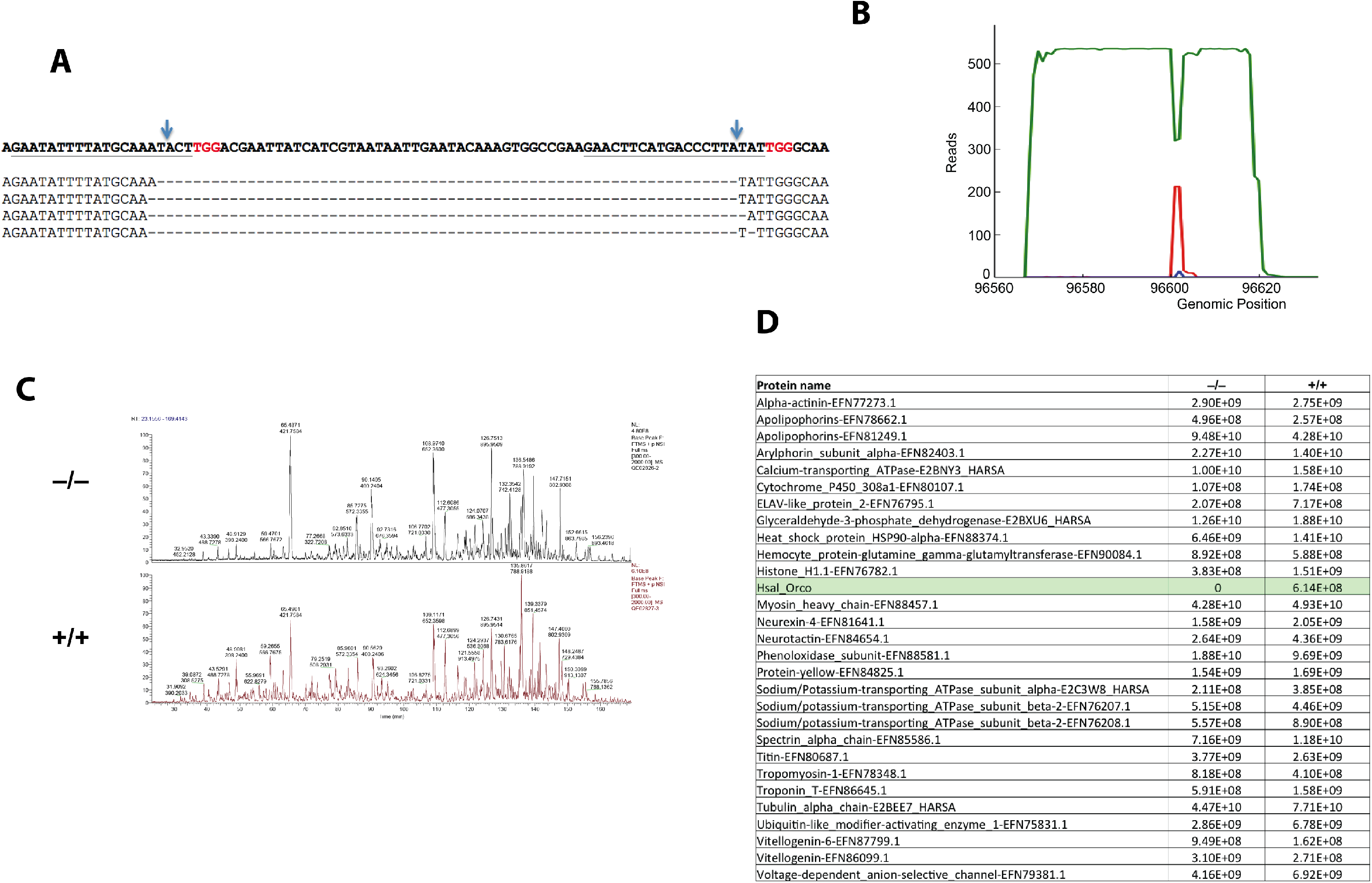
Somatic and germline mutations in *orco*. (A) Somatic mutagenesis with two sgRNAs generated deletions of ~60 nucleotides in length. (B) MiSeq sequencing shows that the somatic mutation rate is approximately 40%. The green line indicates wild type (WT) sequence, while the red and blue lines indicate deletions and point mutations, respectively. Surrounding the target site (genomic position 96003), the WT reads start to decrease, and the mutations start to increase, with the majority of mutations being short deletions. (C,D) Mass spectrometry indicates that no protein could be detected in the antennae of *orco* F4 homozygous mutant. The LC-MS/MS basepeak chromatograms indicate that overall peak profiles are similar in the WT and homozygous female antennae (C). Moreover, the quantifications of peak area in multiple proteins indicate that Orco protein was abundant in the WT antennae but its expression was completely abolished in the homozygote (D).

**Figure S3.**
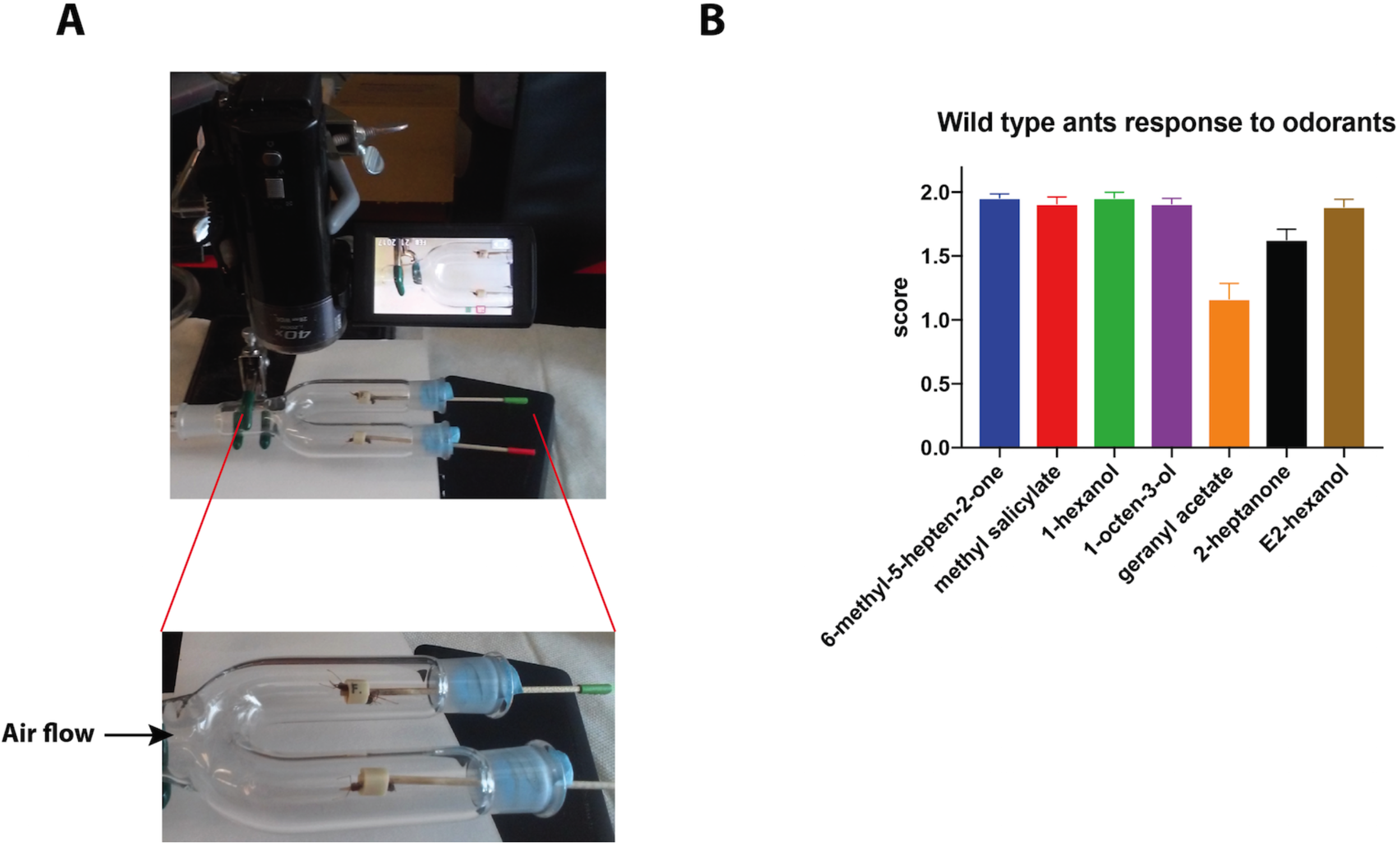
Antennal retraction in response to general odorants. (A) The experimental setting: two ants (one F4 and one WT) were placed in the Y-shape glass tube. Antennal responses to the puff of certain odorants were recorded and scored before knowing the genotypes of F4 ants. (B) Average responses of the 14 wild type ants (3 tests for each ant) to the general odorants (no response: 0; mild response: 1; strong response: 2). Error bars indicate standard error of the mean (SEM).

**Figure S4.**
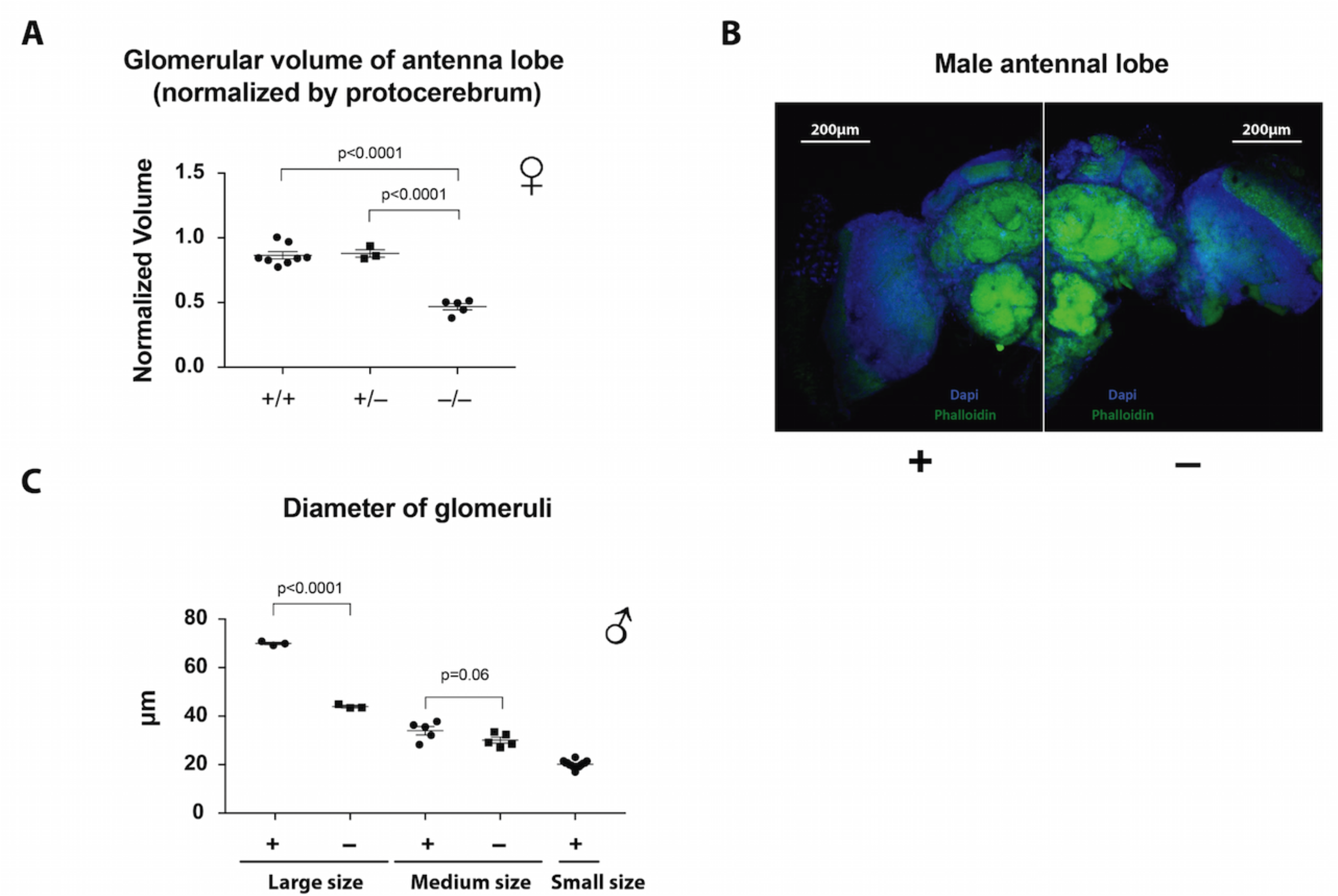
Loss of *orco* in *Harpegnathos* reduced the glomerular volume of antennal lobe and changed glomeruli morphology. (A) Normalized glomerular volume of antennal lobe in WT (n=8) heterozygous (n=3), and homozygous female ants (n=5). (B) The confocal images of wild type (+) vs. mutant males (−): Phalloidin (green), DAPI (blue), 10X objective. (C) The diameter of dorsal glomeruli in WT (n=3): 69.82 ± 0.47 μm and mutant males (n=3): 44.77 ± 0.50 μm. The ventral glomeruli in WT are separated into two groups: medium size (n=5) glomeruli: 34.92 ± 1.72 μm and small size (n=14) glomeruli: 20.5 ± 0.38 μm. There is no statistical significance between medium size glomeruli in WT and mutant ants (n=5): 30.28 ± 1.22 μm, while small glomeruli were not observed in mutant ants. p values are indicated for each plot.

## Reference

Ali, M.F., and Morgan, E.D. (1990). Chemical Communication in Insect Communities-a Guide to Insect Pheromones with Special Emphasis on Social Insects. Biol Rev 65, 227–247.

Benton, R., Sachse, S., Michnick, S.W., and Vosshall, L.B. (2006). Atypical membrane topology and heteromeric function of Drosophila odorant receptors in vivo. PLoS biology 4, e20.

Benton, R., Vannice, K.S., Gomez-Diaz, C., and Vosshall, L.B. (2009). Variant ionotropic glutamate receptors as chemosensory receptors in Drosophila. Cell 136, 149–162.

Blomquist, G.J., Bagnères, A.-G., and MyiLibrary. (2010). Insect hydrocarbons biology, biochemistry, and chemical ecology (Cambridge, UK; New York: Cambridge University Press), pp. 1 online resource (xi, 492 p.).

Bolte, S., and Cordelieres, F.P. (2006). A guided tour into subcellular colocalization analysis in light microscopy. J Microsc-Oxford 224, 213–232.

Bonasio, R. (2012). Emerging topics in epigenetics: ants, brains, and noncoding RNAs. Annals of the New York Academy of Sciences 1260, 14–23.

Bonasio, R., Li, Q., Lian, J., Mutti, N.S., Jin, L., Zhao, H., Zhang, P., Wen, P., Xiang, H., Ding, Y., et al. (2012). Genome-wide and caste-specific DNA methylomes of the ants *Camponotus floridanus* and *Harpegnathos saltator*. Current biology : CB 22, 1755–1764.

Bonasio, R., Zhang, G., Ye, C., Mutti, N.S., Fang, X., Qin, N., Donahue, G., Yang, P., Li, Q., Li, C., et al. (2010). Genomic comparison of the ants *Camponotus floridanus* and *Harpegnathos saltator*. Science 329, 1068–1071.

Boroczky, K., Wada-Katsumataa, A., Batchelor, D., Zhukovskaya, M., and Schal, C. (2013). Insects groom their antennae to enhance olfactory acuity. Proceedings of the National Academy of Sciences of the United States of America 110, 3615–3620.

Capinera, J.L. (2004). Encyclopedia of entomology (Dordrecht; London: Kluwer Academic).

Chiang, A., Priya, R., Ramaswami, M., Vijayraghavan, K., and Rodrigues, V. (2009). Neuronal activity and Wnt signaling act through Gsk3-beta to regulate axonal integrity in mature Drosophila olfactory sensory neurons. Development 136, 1273–1282.

Cong, L., Ran, F.A., Cox, D., Lin, S., Barretto, R., Habib, N., Hsu, P.D., Wu, X., Jiang, W., Marraffini, L.A., et al. (2013). Multiplex genome engineering using CRISPR/Cas systems. Science 339, 819–823.

DeGennaro, M., McBride, C.S., Seeholzer, L., Nakagawa, T., Dennis, E.J., Goldman, C., Jasinskiene, N., James, A.A., and Vosshall, L.B. (2013). orco mutant mosquitoes lose strong preference for humans and are not repelled by volatile DEET. Nature 498, 487–491.

Depickere, S., Fresneau, D., and Deneubourg, J.L. (2004). A basis for spatial and social patterns in ant species: Dynamics and mechanisms of aggregation. J Insect Behav 17, 81–97.

Ditzen, M., Pellegrino, M., and Vosshall, L.B. (2008). Insect odorant receptors are molecular targets of the insect repellent DEET. Science 319, 1838–1842.

Duffield, R.M., Brand, J.M., and Blum, M.S. (1977). 6-Methyl-5-Hepten-2-One in Formica Species-Identification and Function as an Alarm Pheromone (Hymenoptera-Formicidae). Ann Entomol Soc Am 70, 309–310.

Espinosa, J.S., and Stryker, M.P. (2012). Development and plasticity of the primary visual cortex. Neuron 75, 230–249.

Gratz, S.J., Cummings, A.M., Nguyen, J.N., Hamm, D.C., Donohue, L.K., Harrison, M.M., Wildonger, J., and O’Connor-Giles, K.M. (2013). Genome engineering of Drosophila with the CRISPR RNA-guided Cas9 nuclease. Genetics.

Haight, K.L. (2012). Patterns of venom production and temporal polyethism in workers of Jerdon’s jumping ant, Harpegnathos saltator. J Insect Physiol 58, 1568–1574.

Hassan, B.A., and Hiesinger, P.R. (2015). Beyond Molecular Codes: Simple Rules to Wire Complex Brains. Cell 63, 285–291.

Henderson, D.S. (2004). Drosophila cytogenetics protocols (Totowa, N.J.: Humana Press).

Hölldobler, B., and Wilson, E. (2008). The superorganism: the beauty, elegance, and strangeness of insect societies. E W W Norton & Company Inc.

Hölldobler, B., and Wilson, E.O. (1990). The Ants (Cambridge, Mass.: Belknap Press of Harvard University Press).

Hoyer, S.C., Liebig, J., and Rössler, W. (2005). Biogenic amines in the ponerine ant Harpegnathos saltator: serotonin and dopamine immunoreactivity in the brain. Arthropod Struct Dev 34, 429–440.

Hsu, P.D., Lander, E.S., and Zhang, F. (2014). Development and applications of CRISPR-Cas9 for genome engineering. Cell 157, 1262–1278.

Hubel, D.H., and Wiesel, T.N. (1979). Brain mechanisms of vision. Sci Am 241, 150–162.

Jefferis, G.S., Marin, E.C., Stocker, R.F., and Luo, L. (2001). Target neuron prespecification in the olfactory map of Drosophila. Nature 414, 204–208.

Jones, W.D., Cayirlioglu, P., Kadow, I.G., and Vosshall, L.B. (2007). Two chemosensory receptors together mediate carbon dioxide detection in Drosophila. Nature 445, 86–90.

Khila, A., and Abouheif, E. (2009). In situ hybridization on ant ovaries and embryos. Cold Spring Harbor protocols 2009, pdb prot5250.

Kistler, K.E., Vosshall, L.B., and Matthews, B.J. (2015). Genome engineering with CRISPR-Cas9 in the mosquito Aedes aegypti. Cell reports 11, 51–60.

Koutroumpa, F.A., Monsempes, C., Francois, M.C., de Cian, A., Royer, C., Concordet, J.P., and Jacquin-Joly, E. (2016). Heritable genome editing with CRISPR/Cas9 induces anosmia in a crop pest moth. Sci Rep 6, 29620.

Laissue, P.P., and Vosshall, L.B. (2008). The olfactory sensory map in Drosophila. Adv Exp Med Biol 628, 102–114.

Larsson, M.C., Domingos, A.I., Jones, W.D., Chiappe, M.E., Amrein, H., and Vosshall, L.B. (2004). Or83b encodes a broadly expressed odorant receptor essential for Drosophila olfaction. Neuron 43, 703–714.

Li, Y., Zhang, J., Chen, D., Yang, P., Jiang, F., Wang, X., and Kang, L. (2016). CRISPR/Cas9 in locusts: Successful establishment of an olfactory deficiency line by targeting the mutagenesis of an odorant receptor co-receptor (Orco). Insect biochemistry and molecular biology 79, 27–35.

Liebig, J., Hölldobler, B., and Peeters, C. (1998). Are ant workers capable of colony foundation? Naturwissenschaften 85, 133–135.

Liebig, J., Peeters, C., Oldham, N.J., Markstadter, C., and Hölldobler, B. (2000). Are variations in cuticular hydrocarbons of queens and workers a reliable signal of fertility in the ant *Harpegnathos saltator?* Proceedings of the National Academy of Sciences of the United States of America 97, 4124–4131.

McKenzie, S.K., Fetter-Pruneda, I., Ruta, V., and Kronauer, D.J. (2016). Transcriptomics and neuroanatomy of the clonal raider ant implicate an expanded clade of odorant receptors in chemical communication. Proceedings of the National Academy of Sciences of the United States of America 113, 14091–14096.

McKenzie, S.K., Oxley, P.R., and Kronauer, D.J. (2014). Comparative genomics and transcriptomics in ants provide new insights into the evolution and function of odorant binding and chemosensory proteins. BMC genomics 15, 718.

Mombaerts, P. (2006). Axonal wiring in the mouse olfactory system. Annual review of cell and developmental biology 22, 713–737.

Nakagawa, T., Pellegrino, M., Sato, K., Vosshall, L.B., and Touhara, K. (2012). Amino acid residues contributing to function of the heteromeric insect olfactory receptor complex. Plos One 7, e32372.

Oland, L.A., and Tolbert, L.P. (1996). Multiple factors shape development of olfactory glomeruli: insights from an insect model system. J Neurobiol 30, 92–109.

Omasits, U., Ahrens, C.H., Muller, S., and Wollscheid, B. (2014). Protter: interactive protein feature visualization and integration with experimental proteomic data. Bioinformatics 30, 884–886.

Ozaki, M., Wada-Katsumata, A., Fujikawa, K., Iwasaki, M., Yokohari, F., Satoji, Y., Nisimura, T., and Yamaoka, R. (2005). Ant nestmate and non-nestmate discrimination by a chemosensory sensillum. Science 309, 311–314.

Peeters, C., Liebig, J., and Holldobler, B. (2000). Sexual reproduction by both queens and workers in the ponerine ant Harpegnathos saltator. Insect Soc 47, 325–332.

Penick, C.A., Prager, S.S., and Liebig, J. (2012). Juvenile hormone induces queen development in late-stage larvae of the ant *Harpegnathos saltator*. J Insect Physiol 58, 1643–1649.

Perry, M., Kinoshita, M., Saldi, G., Huo, L., Arikawa, K., and Desplan, C. (2016). Molecular logic behind the three-way stochastic choices that expand butterfly colour vision. Nature 535, 280–284.

Petsakou, A., Sapsis, T.P., and Blau, J. (2015). Circadian Rhythms in Rho1 Activity Regulate Neuronal Plasticity and Network Hierarchy. Cell 162, 823–835.

Riabinina, O., Task, D., Marr, E., Lin, C.C., Alford, R., O’Brochta, D.A., and Potter, C.J. (2016). Organization of olfactory centres in the malaria mosquito Anopheles gambiae. Nat Commun 7, 13010.

Robinson, G.E. (1987). Regulation of honey-bee age polyethism by juvenile-hormone. Behav Ecol Sociobiol 20, 329–338.

Sasaki, T., Penick, C.A., Shaffer, Z., Haight, K.L., Pratt, S.C., and Liebig, J. (2016). A Simple Behavioral Model Predicts the Emergence of Complex Animal Hierarchies. Am Nat 187, 765–775.

Schindelin, J., Arganda-Carreras, I., Frise, E., Kaynig, V., Longair, M., Pietzsch, T., Preibisch, S., Rueden, C., Saalfeld, S., Schmid, B., et al. (2012). Fiji: an open-source platform for biological-image analysis. Nature methods 9, 676–682.

Sharma, K.R., Enzmann, B.L., Schmidt, Y., Moore, D., Jones, G.R., Parker, J., Berger, S.L., Reinberg, D., Zwiebel, L.J., Breit, B., et al. (2015). Cuticular Hydrocarbon Pheromones for Social Behavior and Their Coding in the Ant Antenna. Cell reports 12, 1261–1271.

Sperry, R.W. (1963). Chemoaffinity in the Orderly Growth of Nerve Fiber Patterns and Connections. Proceedings of the National Academy of Sciences of the United States of America 50, 703–710.

Suh, H., Deng, W., and Gage, F.H. (2009). Signaling in adult neurogenesis. Annual review of cell and developmental biology 25, 253–275.

Trible, W., Chang, N.-C., Matthews, B.J., McKenzie, S.K., Olivos-Cisneros, L., Oxley, P.R., Saragosti, J., and Kronauer, D.J. (2017). orco mutagenesis causes loss of antennal lobe glomeruli and impaired social behavior in ants. bioRxiv.

Van Oystaeyen, A., Oliveira, R.C., Holman, L., van Zweden, J.S., Romero, C., Oi, C.A., d’Ettorre, P., Khalesi, M., Billen, J., Wackers, F., et al. (2014). Conserved class of queen pheromones stops social insect workers from reproducing. Science 343, 287–290.

Vosshall, L.B., and Hansson, B.S. (2011). A unified nomenclature system for the insect olfactory coreceptor. Chemical senses 36, 497–498.

Vosshall, L.B., Wong, A.M., and Axel, R. (2000). An olfactory sensory map in the fly brain. Cell 102, 147–159.

Watanabe, T., Noji, S., and Mito, T. (2014). Gene knockout by targeted mutagenesis in a hemimetabolous insect, the two-spotted cricket Gryllus bimaculatus, using TALENs. Methods 69, 17–21.

Wicher, D. (2015). Olfactory signaling in insects. Prog Mol Biol Transl Sci 130, 37–54.

Wilt, F.H., and Wessells, N.K. (1967). Methods in developmental biology (New York,: T.Y. Crowell Co.).

Yan, H., Simola, D.F., Bonasio, R., Liebig, J., Berger, S.L., and Reinberg, D. (2014). Eusocial insects as emerging models for behavioural epigenetics. Nature reviews Genetics 15, 677–688.

Yang, B., Fujii, T., Ishikawa, Y., and Matsuo, T. (2016). Targeted mutagenesis of an odorant receptor co-receptor using TALEN in Ostrinia furnacalis. Insect biochemistry and molecular biology 70, 53–59.

Yu, C.R., Power, J., Barnea, G., O’Donnell, S., Brown, H.E., Osborne, J., Axel, R., and Gogos, J.A. (2004). Spontaneous neural activity is required for the establishment and maintenance of the olfactory sensory map. Neuron 42, 553–566.

Zhao, N., Guan, J., Ferrer, J.L., Engle, N., Chern, M., Ronald, P., Tschaplinski, T.J., and Chen, F. (2010). Biosynthesis and emission of insect-induced methyl salicylate and methyl benzoate from rice. Plant Physiol Biochem 48, 279–287.

Zhou, X., Rokas, A., Berger, S.L., Liebig, J., Ray, A., and Zwiebel, L.J. (2015). Chemoreceptor Evolution in Hymenoptera and Its Implications for the Evolution of Eusociality. Genome biology and evolution 7, 2407–2416.

Zhou, X., Slone, J.D., Rokas, A., Berger, S.L., Liebig, J., Ray, A., Reinberg, D., and Zwiebel, L.J. (2012). Phylogenetic and transcriptomic analysis of chemosensory receptors in a pair of divergent ant species reveals sex-specific signatures of odor coding. PLoS genetics 8, e1002930.

Zube, C., and Rössler, W. (2008). Caste- and sex-specific adaptations within the olfactory pathway in the brain of the ant Camponotus floridanus. Arthropod Struct Dev 37, 469–479.

